# How does perception of zoo animal welfare influence public attitudes, experiences, and behavioural intentions? A mixed-methods systematic review

**DOI:** 10.1101/2024.03.20.585889

**Authors:** Nicki Phillips, Laëtitia Maréchal, Beth Ventura, Jonathan Cooper

**Author notes:** Corresponding authors (NP), (LM).

## Abstract

The public expects zoos to provide high standards of animal care. Failing to meet public expectations can have detrimental impacts on public experiences and behaviour, which in turn can compromise zoos’ organisational goals relative to conservation and public education. Despite increased research interest in understanding how the public perceives animal welfare in zoo settings, to date the factors that influence such perceptions are still unclear. To address this gap in knowledge, we conducted a mixed methods systematic review using a PRISMA approach to identify the factors that influence public perceptions of zoo animal welfare and the potential ramifications of these perceptions on public attitudes, experiences, and behaviours. A total of 115 peer reviewed journal articles were analysed: 43 provided qualitative data for thematic synthesis and 85 reported quantitative data for content analysis. Three main groupings were identified that impacted public perception of animal welfare in zoos: human, animal, and environmental factors. Within the human factors, ethical justifications, direct interactions, and inappropriate visitor behaviours were important. For the animal factors, animals’ behaviour, apparent health status, and the suitability of certain taxa for captivity were found to be key. Finally, several aspects of the environment -- conditions of the facility, the exhibit, and welfare-related educational material --were influential. Overall, negative perceptions of animal welfare resulted in negative visitor attitudes towards zoos, detrimentally impacted experiences, and lowered likelihood to visit zoos and engagement in conservation efforts. The articles in this review provided valuable insights into the factors affecting public perception of zoo animal welfare; however, future research may benefit from a more structured approach to increase comparability and validity of results across studies. We conclude by proposing seven recommendations to increase the robustness and validity of future research in this area.

## 1.0 Introduction

Public concern for the welfare of captive animals has been increasing and high standards of care are expected across a range of captive settings [1]. These expectations extend to all zoo facilities, including wildlife and safari parks, aquariums, and sanctuaries. When care standards are perceived to be insufficient, poor visitor perceptions of zoo animal welfare can occur, which in turn can reduce financial support from the public and engagement with conservation initiatives [2]. Consequently, reduced public support can negatively impact the ability of zoos to achieve organisational goals to achieve high animal welfare, effective public education, and conserve species [3]. Considering the potential consequences of negative zoo visitor perceptions, understanding how animal-visitor interactions impact visitors is important. Much of the research regarding animal-visitor interactions in zoos has primarily focused on the impact of those interactions on the animals [4], with fewer studies on impacts on zoo visitors [5]. Several literature reviews have provided valuable contributions towards understanding visitor experiences at zoos [3–5, 7, 8], but to our knowledge no systematic review has been undertaken. More specifically, we presently lack a robust understanding of the impacts of zoo animal welfare perceptions on visitor experiences and behaviours, in addition to the specific factors influencing such perceptions.

Positive and negative perceptions of animal welfare have the potential to influence visitor behaviour. For example, the likelihood that the public will visit or re-visit a zoo may be reduced if negative welfare is perceived [2], though negative welfare perception is not always a barrier to zoo attendance [9]. Various justifications for continued visiting when welfare concerns are evident have been supplied by visitors in attempts to reduce cognitive dissonance [10]. Supporting conservation goals appears to be a common justification [11]; for instance, Curtin and Wilkes [10] found participants in swim-with-dolphin encounters used the perceived conservation activities of these facilities to reduce dissonance when welfare concerns became evident. In contrast, the likelihood of visitors providing conservation donation support may be reduced due to negative welfare perceptions [2]. As good welfare, visitor recreation, and conservation are common objectives of modern zoos [12], and as each objective can potentially be impacted by visitor perceptions of zoo animal welfare, understanding why these attributes influence perceptions is crucial.

Several factors (hereafter referred to as zoo attributes and used in this review to describe variables representing human, animal, and environmental-level factors affecting zoo visitor perceptions and experiences) have been highlighted as influential to public perception of zoo animal welfare [5]. Human, animal, and environmental zoo attributes mirror those described in farmed (reviewed by [13]) and zoo animal welfare measurement [14], and in the One Welfare concept, which “*recognises the interconnections between animal welfare, human wellbeing, and the environment*” [15]. For example, for human-level factors, inappropriate visitor behaviours such as feeding or teasing animals can detrimentally impact visitor perceptions of zoo animal welfare [11]. At the animal level, behaviour perceived to be undesirable or unnatural, such as stereotypic pacing, tends to diminish visitor perceptions of zoo animal welfare [2], while perceiving animals as engaging in play behaviours generally translates to perceptions of good welfare [16]. Additionally, visitor misinterpretation of animal behaviour (e.g. inactivity perceived as abnormal in animals who naturally sleep for large portions of the day) can reduce welfare perceptions [16]. With respect to environment, naturalistic exhibits (i.e. those designed to mimic aspects of an animal’s natural habitat) may aid in improving visitor perceptions of zoo animals and zoo animal welfare [17], as does provision of environmental enrichment [18]. While there are some indications that welfare-related interpretation may enhance zoo visitor perceptions [19], research is currently limited.

Research investigating how the public perceives animal welfare in zoo settings has increased, yet the factors which are influential to such perceptions remain unclear. Visitors have also shown conflicting perceptions of the same factor when considering implications to welfare. As perceptions of zoo animal welfare may impact visitor behaviour, a comprehensive review focusing specifically on zoo animal welfare perceptions and the impacts of these perceptions on visitor experience and behaviour is needed. Hence, this systematic review aims to build on the foundations supplied by previous research, with the following objectives: 1) to identify and analyse existing literature exploring what zoo attributes, i.e. animal, human and environment, influence visitor perceptions of animal welfare, either positively or negatively, and 2) to characterise how positive or negative perception of animal welfare impacts visitor attitudes, experiences, and behaviour. This review was conducted following The Preferred Reporting Items for Systematic Reviews and Meta-Analysis (PRISMA) 2020 Guidelines [20] (Supplementary material S1).

## 2.0 Materials and Methods

The review began with a search of existing scientific, peer-reviewed literature. The search strategy (Supplementary material S2) was assessed for appropriateness by an academic librarian, and searches were first executed on Google Scholar, SCOPUS, Web of Science and EBSCOhost in June 2022, and repeated in August 2023 to capture additional literature published in the time since the review was initiated. These searches resulted in a combined 115 articles included in this review (Fig 1). A full and independent search and screening process was conducted by two researchers (NP and BV) to validate results. The review protocol was pre-registered on Open Science Framework under the registration number: hc6qx (Supplementary material S3).

**Fig 1.**
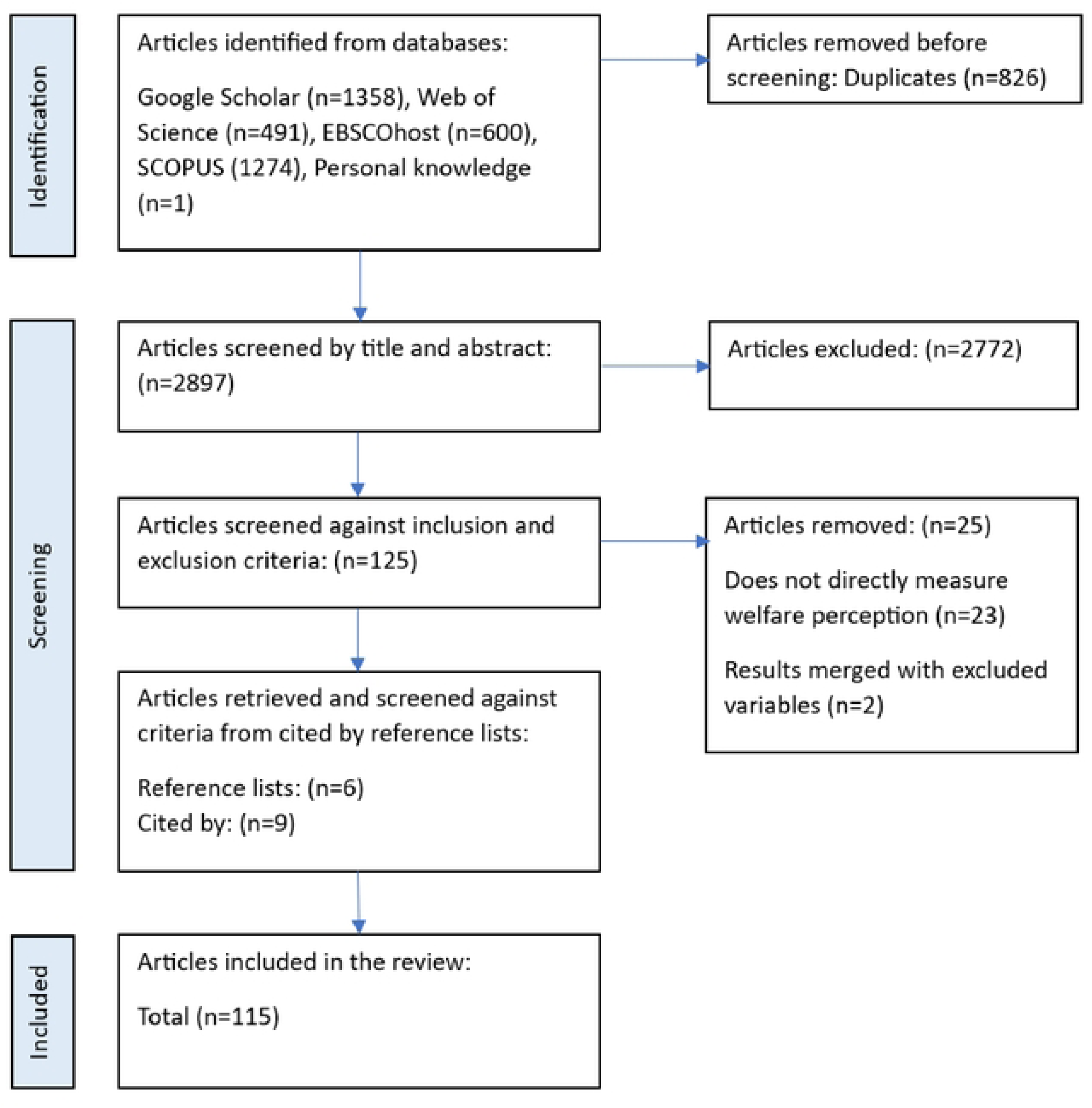
PRISMA flow diagram displaying the search and assessment process which resulted in final inclusion of 115 articles in this systematic review.

### 2.1 Screening

Studies met inclusion criteria if they directly measured zoo animal welfare perception and/or evaluated the impact of welfare perception on visitor attitude, experience, and intended or actual behaviour. Animal welfare was defined as by Broom [21]: “*the welfare of an individual is its state as regards its attempts to cope with its environment.”* We recognise there are several animal welfare frameworks in the literature. Here, we rely on the Five Domains Model [22], which recognizes that animal welfare is affected by nutrition, physical environment, health, behavioural interactions, and mental state. Included studies could be quantitative, qualitative, or mixed methods, take place within or outside of zoo facilities, and be based in any geographical location. Participants could be any age. Studies were excluded if they were not published within a peer reviewed journal, or if full text was not available or not available in the English language. ‘Grey literature’ (e.g. book chapters, unpublished works, conference papers/proceedings that are not in peer reviewed journals, theses and dissertations, reports, newsletters) were also excluded. Studies published before 1990 were excluded as significant changes to zoo husbandry practices and conservation initiatives occurred around this time [23], making direct comparisons to earlier studies difficult. Participants could not be associated with zoos professionally. Participants could be either visitors to a zoo facility at the time of study participation, or members of the public at a remote location. Perceptions must have clearly and directly related to zoo animal welfare; for example, participant opinion of a zoo attribute (such as enrichment items) would not be included in the absence of a clear link between opinion and perceived impacts to zoo animal welfare. Zoos and zoo animals were defined as by the licencing requirements laid out in the UK *Zoo Licensing Act 1981* [24]: *“ any establishment where wild animals (as defined by section 21) are kept for exhibition to the public, otherwise than for purposes of a circus (as so defined) and otherwise than in a pet shop” and “the public have access, with or without charge for admission, on seven days or more in any period of twelve consecutive months”.* The term ‘wild animals’ is defined by ZLA 1981 as “*“animals of the classes Mammalia, Aves, Reptilia, Amphibia, Pisces and Insecta and any other multi cellular organism that is not a plant or a fungus and “wild animals” means animals not normally domesticated in Great Britain”.* Therefore, ‘wild animals’ are defined by this study as those not normally domesticated in the zoo’s geographical region. Elephant camps and tiger temples in Southeast Asia were excluded as they fell outside of the scope of this review. See Supplementary material (S4) for articles excluded at this stage.

### 2.2 Data extraction and quality assessment

An inclusive approach (as described by [25]) was followed wherein all text relating to the review question within the findings/results sections was included, alongside any relevant text found within abstracts and discussions. If articles discussed multiple facilities including those other than zoos, only content directly and clearly relating to zoos were extracted (n=11 articles). When articles shared data sets, only unique observations/results from each paper were extracted (n=8 articles). The Mixed Methods Appraisal Tool (MMAT) version 2018 [26] was used for quality assessment (Supplementary material S5) to inform analysis and interpretation of results, not to remove studies perceived to be poor.

### 2.3 Analysis

As included papers presented both qualitative and quantitative data, our data analysis was comprised of two complementary analyses: a thematic analysis (qualitative data) and content analysis (quantitative data). Analysis began with an inductive thematic synthesis of qualitative data (as described by [25]), using NVivo 12 software (NVivo 12, QSR international, London, UK). The process began with familiarisation of the data via multiple readings, followed by coding into descriptive themes. This was a cyclical process as more influential factors to zoo animal welfare perception became apparent and the themes required refinement. Once the broad themes had been identified, coding was repeated in a hierarchical tree structure to develop analytical themes, which provided a more in-depth understanding of the influential factors to zoo animal welfare perception and the potential impacts to visitor behavioural intentions. This step involved identifying what factors were influential, but also why these factors were influential, either positively or negatively.

The themes and subthemes identified from the qualitative analysis then informed the coding scheme used to process the quantitative data as follows: content analysis (as described by [27] was performed. This process also began with familiarisation of the data. Familiarisation was followed by coding the data into the themes and subthemes identified in the thematic synthesis, whilst remaining alert to any potential new themes. All the themes and subthemes created during the thematic synthesis were also evident in the quantitative data, except the subtheme facility size which was exclusive to the qualitative data. Finally, all the studies (both qualitative and quantitative) were re-read to ensure no theme relevant to the research question was overlooked within the extracted data. By including both quantitative and qualitative data in a convergent integrated approach, a more comprehensive understanding of visitor perceptions could be gained. Each data set was synthesised narratively [28] which provided a textual description and analysis of combined findings. Repeat assessment of themes occurred with consideration of articles highlighted as containing potential risks of bias (Section 2.2), to ensure the validity of the evidence and therefore the validity of the themes created.

## 3.0 Results and Discussion

### 3.1 Descriptive summary of included articles

Due to the similarity of the results obtained from the thematic and content analyses, results are presented together. A total of 115 articles were included in the review, with 43 supplying qualitative data for thematic synthesis and 85 supplying quantitative data for content analysis (some articles supplied both qualitative and quantitative data). Articles spanned 25 years (1993-2023), with the highest rate of articles published in 2021 (14.8% of total). Research populations were mainly based in North America (30.8%), followed by Asia (17.7%), the United Kingdom (16.9%), Europe (13.8%), Oceania (10.8%), Africa (7.7%) and South America (2.3%). 60 journals were represented, with most articles published in Zoo Biology (12.2%) or Anthrozoös (12.2%), Animals (6.1%), the Journal of Zoo and Aquarium Research (5.2%), and Visitor Studies (4.3%). Five classes of animal were represented, with most studies including mammals (56.7%), 8.5% including birds or reptiles, and only 1.4% including amphibians or insects (Figure 2). Within the class Mammalia, the most represented orders were Carnivora (34.2%), Primates (29.1%), and Artiodactyla (even-toed ungulates, 8.9%). Most research focused on facilities described as zoos (76.9%), with 6.7% relating to aquariums, 5.2% to marine parks, 3.7% safari parks, 3.0% sanctuaries and wildlife parks and 1.5% not stating a facility (i.e. while taking place in a zoo, the specific zoo name and facility type were not supplied). A full list of all included studies and their relative theme contribution can be found in the Supplementary material (S6).

**Fig 2.**
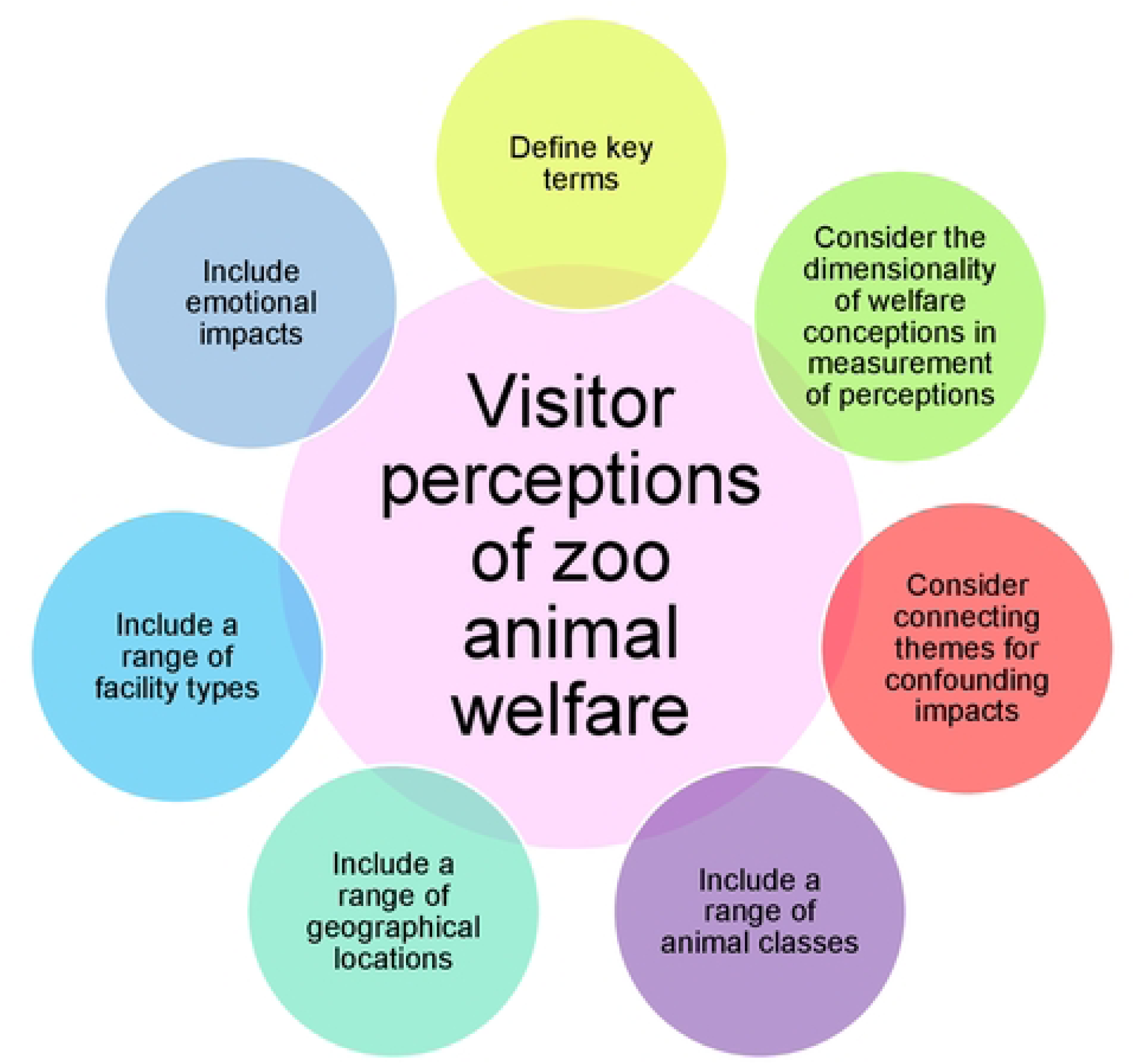
Recommendations for the study of visitor perceptions of zoo animal welfare.

25.2% of the total articles assessed visitor perceptions of welfare via perceived affective state, 20.8% of articles via health indicators, 13.9% via natural behaviour display and 17.3% via perceived quality of care. 38.2% of the total articles used a statement of perceived general overall welfare, while 33.9% of articles did not use a specific category but asked open questions, leading to conceptions or descriptions of welfare perceptions based on participants’ own words. The five domains model (Mellor) was not used to categorise the types of welfare perception assessments, as perceptions such as quality of care and those based on participants’ descriptions did not meet the criteria for any specific domain. 65.2% of the total articles used one category of welfare perception (i.e. perceived affective state, health, natural behaviour, quality of care, or overall welfare perception), 52% of which utilised open questions and participant wording. 22.6% of the total articles utilised two categories of perception, 10.4% used three, and 1.7% used four.

### 3.2. Overview of themes

Identified themes related to objective 1 (i.e. influential factors to welfare perceptions) were organised within three main categories: human-, animal-, and environmental-related factors. Three themes were described within human-related factors affecting perceptions toward zoo animal welfare (Table 1): ethical considerations (with subthemes of beliefs about captivity, and animal rights); direct human-animal interaction; and inappropriate visitor behaviour. The term ‘direct human-animal interactions’ denotes interactions between humans and animals during encounters, shows, training exercises, and husbandry practices. The animal category consisted of three themes (Table 1): disproportionate suffering, animal behaviour, and apparent health status. The environmental category included three themes (Table 1): the facility (with subthemes of zoo purpose and advertisement, and size), the exhibit (with subthemes of naturalistic enclosure, exhibit size, enrichment, group size, condition of the enclosure, diet, sensory stressors, and temperature), and welfare interpretation. One theme was described which addressed objective 2 (i.e. potential impacts of welfare perceptions on visitor experience and behaviour): impacts of welfare perceptions on visitor experience and behaviour. Themes were categorised as linked when one attribute highlighted how and/or why the other attribute was influential to welfare perceptions (Table 1). For example, large enclosure size was perceived to facilitate natural behaviour, which was perceived to indicate good welfare. See Supplementary Information (S7) for a breakdown of each theme (e.g. number of studies in which themes appear and which animal orders and type of zoo facility were represented). Visitor perceptions of zoo animal welfare begin to form before visits have begun, due to variations in preconceived beliefs concerning captivity [29]. Thus, human-level factors regarding ethical perceptions of captivity are the opening themes of the discussion (below).

**Table 1.**
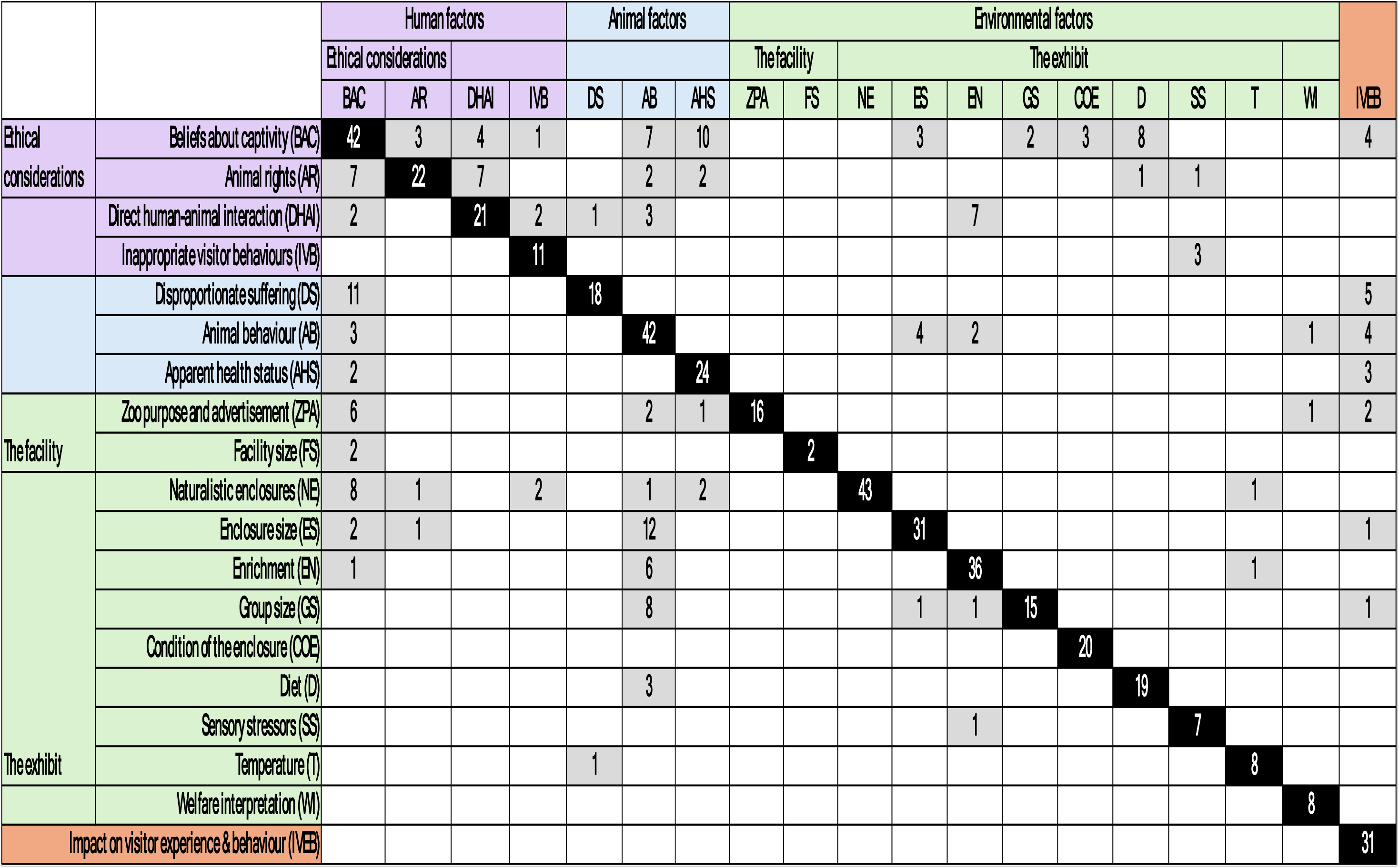
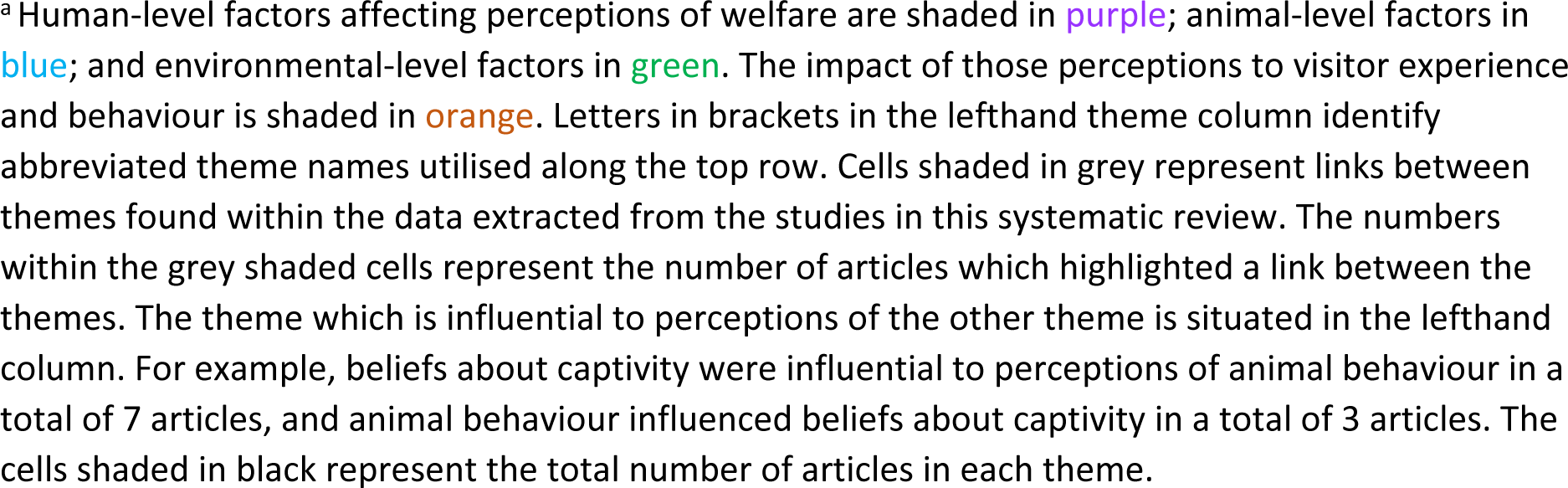
Themes identified as affecting perceptions of zoo animal welfare and impacts of those perceptions on visitor experience and behaviour.

### 3.3 Human-level factors

#### Ethical considerations

##### Beliefs about captivity

Preconceived beliefs regarding captivity were identified as a factor impacting perceptions of zoo animal welfare, as zoos were perceived to be protectors of vulnerable animals and providers of quality care [9–11, 18, 29–48]. For example, sentiments expressed included, “*they have protection and a good food supply*” [39]. Origins of the animals in zoo collections impacted perceptions of zoos as protectors, as ‘rescued’ or captive-bred animals were deemed more acceptable than wild caught [9–11, 43, 44]. Captive breeding was perceived to habituate animals to the captive environment and therefore reduce suffering [10, 33, 34, 39] and the provision of care and/or protection sometimes led to the belief that some animals were better off in captivity than in the wild [9, 11, 35, 41, 43, 44]. Holding such views appears to attenuate otherwise uneasy feelings about the ethics of captivity, as described by Curtin and Wilkes [10]: *“several respondents attempted to reduce dissonance by emphasising that the dolphins were either bred in captivity and therefore had become accustomed to it or had been rescued”*.

The release of animals into the wild --either after they have recovered from the circumstances they were rescued from or through participation in breeding programmes-- was also considered important to ethical acceptance of zoos [11, 32, 39], highlighting that visitors want zoos populated by endangered and/or rescued animals who will ultimately be released into the wild. Yet, zoos house many species who are neither endangered, rescued, or part of release programmes [49]; furthermore, two studies documented assumptions by visitors that zoos were releasing animals into the wild, without the facilities making any such claim [50, 51]. Reasons behind this mistaken assumption were unclear.

In contrast, captivity was perceived to negatively impact zoo animal welfare more often than positively impact welfare in this theme [9–11, 16, 18, 29, 31–39, 41, 45, 46, 52–67]. Articles highlighted beliefs amongst visitors that zoo animals are wild animals imprisoned and denied the freedom of their natural habitats, which zoos can never replicate. Failure to replicate nature was perceived to restrict natural behaviours and lead to boredom, stress, unhappiness amongst animals, and in some cases, this belief was linked to strongly negative perceptions of zoos. For instance, some participants were said to “*perceive zoos as places where animals are often abused, suppressed and congested in limited/small spaces”* [29].

In addition to protection and provisions of care [33, 43, 44], several additional justifications were supplied in the ethical defence of zoos; these included education and research opportunities [9–11, 32–34, 38, 43, 44, 68], conservation [10, 11, 29, 34, 36, 43, 44], human safety (i.e. safer in comparison to viewing animals in their natural habitat and also increasing safety by removing and preventing the return of animals from natural environments shared by humans and animals) [11, 33, 39, 43, 44], and entertainment [9, 29, 33, 34, 36, 41, 43, 44, 67, 68]. Collectively, benefits to humans were considered as much as, or more often, than benefits to animal welfare when justifying captivity. Although, all animals were perceived to benefit from the few, even if poor welfare was perceived as they were viewed as “*taking one for the team”* [34] and giving people *“a more in depth understanding of why animals should not be contained”* [9]. This perceived sacrifice was viewed as preventing the need for others of their kind to suffer captivity in their place, or potentially in the future.

##### Animal rights

Study participants believed that zoo animals have a right to life [48], to appropriate end-of-life care [69], to choice and control over their environments; consequently, provision of these rights was integral to perceptions of animal welfare as good or bad. Beliefs involving animals’ freedom of choice included access to preferred food items [39], chosen activities such as swimming or playing [16, 41] (as animals should be able to “*do whatever they want”* [41]), and ability to remain within private areas unseen by visitors [38, 46, 47, 54, 55, 70] and away from noise [50]. Furthermore, an animal’s right to choose to participate in direct interactions including training, performance activities, and receiving direct human contact was highlighted as important to welfare and thus linked to ethical acceptability of these practices [10, 11, 36, 71, 72]. This view was captured by one participant who shared: “*it’s one thing if the animal wants to do it, and it’s a different thing if you stab him with a stick”* [11]. Perceptions that animal rights were discarded or suppressed by zoos led to feelings of ethical unease and dislike of zoos [29, 32].

#### Direct human-animal interactions

The participation of animals in encounters and shows was often perceived as unnatural and thus potentially detrimental, both by visitors taking part in these encounters and by general visitor populations [10, 32, 36, 42, 52, 73]. Direct or close physical contact with humans was highlighted as particularly concerning [10, 37, 71, 74, 75], though this was not the consensus view of study participants. Conversely, direct/close contact was frequently perceived positively as an indication of a close bond between trainers/zookeepers and the animals, which increased perceptions of quality care and welfare [9, 36, 46, 62, 72, 73, 76]. Perceptions of welfare as impacted by keeper-animal contact were mediated by species [72], and by staff qualification level [43, 44, 73]. Similar contrasts were seen in perceptions of welfare as influenced by training, with training perceived to provide physical and psychological enrichment in which animals enjoy participating [10, 36, 42, 62, 73, 77], but alternatively being stressful, unnatural, and demeaning [10, 32, 36, 42, 52, 73]. Improved views toward training were more frequent if it was perceived to avoid aversive training approaches and instead rely on rewards [11, 43, 44, 70]; illustrated by comments like, ““*you don’t need to use a whip, and you don’t need to use something that is not positive reinforcement”* [11]. Whether an animal was seen as able to choose to participate in training or other forms of direct contact with humans was important to perceptions of animal enjoyment [9–11, 36, 52].

#### Inappropriate visitor behaviour

Other visitors’ behaviours were perceived to impact zoo animal welfare [43, 44]. Behaviours perceived to negatively impact welfare included: feeding animals [11, 32, 51, 68, 70, 78]; touching animals when they were not supposed to [51, 70] or not following guidelines when doing so [11, 71]; using flash photography in aquariums (e.g. perceived to “*scare the little creatures”* [71]), making too much noise [32, 68, 79]; teasing [11, 78, 80]; and throwing items at animals [11, 68]. Informing visitors of which behaviours can be detrimental to welfare and requesting a reduction or end to these behaviours may therefore not only improve the welfare of the animals, but also improve the welfare perceptions of other visitors. However, research exploring the efficacy of this form of interpretation is conflicting [81–87]. Further work is suggested to inform approaches to promoting appropriate visitor behaviour, including more efficacious methods of welfare interpretation.

### 3.4 Animal-level factors

#### Disproportionate suffering

Zoos’ perceived (in)ability to provide appropriate habitats impacted the acceptability of captivity and thus welfare perceptions toward several taxa, including cetaceans [9, 10, 36, 56, 67], polar bears [41], primates [57, 58, 60], elephants [46, 88, 89], birds [11, 66], sloths [30], and pythons [30]. Perceived intelligence [10, 46], sentience [60], and large size [90] were also contributing factors to perceived negative welfare. Explanations included, “*the more intelligent I think an animal is, the more I think they are prone to boredom and that is what puts me off zoos or other captive displays. You can see that they get incredibly bored and the more intelligent the creature, the worse it is"* [10]. Even in the face of adversity (i.e. climate change), participants *"emphasized the fact that a polar bear is part of nature and belongs there, despite the fact that a polar bear in the wild needs to work hard for survival. This group also believed that the polar bear actually does not belong in a zoo"* [41]. Yet, when members of the public were surveyed to explore the most desired traits of zoo animals, and which animals they most desired seeing, several species with these same traits (e.g. charismatic megafauna such as gorillas) were reported as desirable [91]. This creates a potential conflict for zoos, who need to house enough desirable species to attract visitors while simultaneously providing reassurance that these animals –possessing traits which also make them more challenging to house in welfare-friendly ways-- receive adequate care within a captive setting. Indeed, a noted influential factor to captivity acceptability was ‘likeability’, as more ‘likable’ animals also garnered higher levels of welfare concern [36, 92, 93].

#### Animal behaviour

Many studies found that behaviours which visitors perceived to be active and natural were considered positive indicators of welfare [10, 16, 19, 35, 38, 41, 43, 46, 52, 54, 59, 62, 70, 77, 88, 94–105]. In contrast, behaviours perceived as abnormal or inactive were thought to be negative indicators [2, 16, 19, 38, 40, 41, 47, 61, 64, 71, 73, 95, 99, 104, 106–108]. For example, inactivity was perceived negatively as “*they lay around depressed or sit as if bored”* [16]. Yet, visitors may not always accurately judge the so-called ‘naturalness’ of behaviours [104], nor may they consistently equate natural behaviours with good welfare [95].

Additionally, not all natural, active behaviours were perceived positively by zoo visitors, as several studies found that non-stereotypic walking behaviour was viewed as a neutral [95] or negative welfare indicator [61, 105]. Visitors may have perceived this locomotion to be abnormal, as walking perceived to be stereotypic pacing was often thought to indicate compromised welfare [2, 64, 104, 108]; however, differentiation between stereotypic pacing and normal active behaviour appeared to be difficult for many visitors [19, 104]. Therefore, several authors have stated the need for further research to explore visitor perceptions of stereotypic behaviour and the impacts of interpretation on these perceptions [2, 3, 5, 104]. We recommend that future research includes attention to how visitors categorise observed behaviours (natural/abnormal). More broadly, given the general lack of research on visitor understanding and perception of stereotypic or other abnormal behaviours (e.g. excessive grooming), a better understanding of visitor accuracy regarding identification of stereotypic behaviours and their influence on perceptions may help inform more effective interpretation in this area.

There appears to be comparably less research focused on visitors’ interpretation of and perception toward social behaviours and relationships between animals as affecting their welfare, and what exists appears conflicting. For example, ‘timidness’ and aggression were sometimes perceived to indicate negative welfare [53, 98], yet displays of dominance were also thought to indicate positive welfare [16]. Impact of perceived aggression on welfare perceptions may vary amongst species, as for example Markwell [109] reported that zoo visitors expected to witness aggression in Tasmanian devils and were disappointed if aggressive behaviour was absent. More consistently, beliefs about play and affiliative social behaviours (like allogrooming), translated to perceptions of positive welfare [10, 16, 36, 47, 72, 73, 95, 96, 99, 110, 111]. Yet play behaviours can appear extreme and even violent [112], and communication signals mediating social interactions between animals are often quite subtle [113]. Whether visitors can accurately identify the function of such behaviours is therefore expected to mediate welfare perceptions, but this requires further research.

Finally, opportunities for breeding were considered beneficial to welfare [38, 54, 72]. However, not all animals are given this opportunity, and when the opportunity does arise, the repercussions are not always acceptable to the public. For example, breeding may contribute to surplus animals and euthanasia, which can potentially result in decreased visit likelihood [114].

#### Apparent health status

Several studies have highlighted that the provision of appropriate and adequate medical care is expected and considered a requirement of good welfare for zoo-housed animals [11, 38, 43, 44, 55, 68, 100]. Perceptions of good health were evident when animals appeared “*well groomed*” [16], as this signified that animals “*were respected*” [48]. Additionally, signs of a healthy weight were viewed positively [16], but negative perceptions arose when animals were perceived to show signs of injury or disease, appeared in poor physical condition, or were under or overweight [16, 19, 29, 54, 59, 64]. Indeed, expectations for robust health may override other expectations: for example, animals may have large, naturalistic enclosures with ample enrichment, but if they appear to be injured or unwell, perceived poor health status may be an overriding factor [115].

The degree to which physical health was prioritised against other welfare issues varied among studies. For example, Packer et al. [16] found prioritisation of health over affective state and perceived quality of care, while Warsaw and Sayers [19] documented greater emphasis placed on behavioural needs and naturalistic environments over health by zoo visitors. In other cases, no significant differences were found [54]. Moreover, perceptions of zoo animal health were influential to visitor experience in some cases [16, 98], but not others [107], and perceptions of animal happiness were more important to some visitors’ emotional connections than health [16]. The inclusion of multiple welfare dimensions in future research, including health, may generate a more holistic picture of visitor perceptions of zoo animal welfare and potential impacts to visitor experiences and behaviours.

### 3.5 Environmental-level factors

#### The facility

##### Zoo purpose and advertisement

How zoos identify themselves can shape public perceptions of welfare, with wildlife and safari parks favoured over zoos [43, 44], and marine parks viewed more critically (e.g. perceived as abusive [9, 56, 67] and lacking in educational value [9]). Zoos who promote themselves as centres for education and conservation were perceived to provide higher welfare than those advertised primarily as places for entertainment [29, 31, 34]. A common objective advertised by zoos is research [12], though terminology (specifically the term ‘scientific research’) can detrimentally impact perception of welfare by increasing perceived negative impacts of captivity and decreasing the perceived importance of zoo animal-based research [53]. Although considered ethically justifiable (See *beliefs about captivity*), research was sometimes viewed as unenjoyable for the animals taking part [116]. However, this trend was inconsistent, as other work found strong visitor agreement with the importance of conducting research in zoos and beliefs that research was good for the animals [117]. Additionally, animals in the wild were also perceived to benefit from research, for example, “*the research that’s done there allows biologists to know more about how to keep them happy and healthy…. And all that research hopefully will be able to increase they longevity and livelihood in their natural habitat”* [94].

Images used in zoo advertising also impact perceptions, with human proximity to animals [118] and keeper presence and captive settings [63] generally resulting in more negative welfare perceptions, though effects varied by species and not all studies have found this effect [30]. Interestingly, negative perceptions associated with zoo images increased the expressed likelihood of conservation donations, though this also varied between species [63].

##### Facility size

Relatively few studies have assessed the impact of establishment size on welfare perceptions, but there is some suggestion that smaller establishments were viewed as less commercial and exploitative [29], but also less governed by adequate welfare laws and lacking in sufficient resources to provide quality care [11].

#### The exhibit

##### Naturalistic enclosures

Consistent amongst the reviewed articles was that naturalistic exhibits (i.e. exhibits designed to be aesthetically appealing and mimic natural habitats [58]) were associated with more positive welfare perceptions [11, 16, 18, 19, 29, 30, 32, 35, 37, 39, 40, 42–44, 46, 47, 51, 52, 62, 64, 65, 70, 88, 95–98, 105, 109, 119–129]. However, naturalistic enclosures may impinge on expression of natural behaviour as naturalistic aesthetics do not always equate to usable space [95, 130]. Even if the enclosure is naturalistic, the presence of barriers was perceived negatively if the materials used were reminiscent of bars and/or cages as animals were perceived as confined and imprisoned [29, 39, 47, 52, 64, 65], even where the use of these barriers can increase the animals’ useable space [130]. However, enclosures perceived to allow for (apparent) free ranging were perceived as beneficial to welfare [32, 51, 124], even if these same enclosures exposed animals to perceived risk of harm [32, 51, 71]. Ultimately, how visitors weigh the relative benefits of so-called natural environments against behaviour expression is unclear, suggesting the need for further research [95].

##### Enclosure size

Unsurprisingly, enclosures perceived as small were consistently associated with more negative welfare perceptions, compared to larger, apparently spacious, enclosures which were perceived to better meet animals’ needs by providing adequate space for natural behaviours and physical comfort [10, 11, 16, 29, 34, 35, 40, 41, 43, 44, 46, 47, 51, 52, 54, 55, 59, 62, 64, 70, 88, 90, 97, 98, 109, 111, 120, 128, 131]. How big an enclosure should be (for each species) and how visitors render judgement remains ambiguous, as qualitative findings have often been limited in their description (e.g. *“looked too small”* [40]). Perceptions of enclosure size adequacy may be influenced by group size [132] and exhibit location (indoors potentially worse than outdoors) [90], but visitor rationale for these judgements also remains unclear. Furthermore, enclosure size perceptions may be influenced by animal behaviour, with perceived abnormal behaviour [64] and natural behaviour restriction [10, 47] reducing size adequacy perceptions, and perceived natural behaviour display increasing perceptions [16, 41, 55].

##### Enrichment

Use of environmental enrichment consistently had a positive impact on welfare perceptions [18, 29, 35, 37, 38, 43, 44, 47, 50, 54, 62, 64, 73, 90, 93, 94, 96, 97, 99–102, 107, 110, 116, 119, 131, 133–139], including feeding-related enrichment [99–102, 136, 138]. Positive perceptions of feeding enrichment are not surprising, as the act of feeding in any circumstance increased welfare perceptions [61, 95, 99, 102]. Yet not all visitors were aware of enrichment in general [139] or which items within an enclosure are enriching [40, 99]. While both naturalistic and unnaturalistic enrichment were perceived positively [40, 99, 102, 119, 134, 135], unnaturalistic enrichment may be more easily identifiable to visitors [40]. Visitor perceptions may be enhanced or improved by the knowledge of enrichment strategies [18]; more targeted signposting as to enrichment strategies in use within zoo enclosures may be useful in improving visitor knowledge of enrichment.

##### Group size

Many studies identified the importance of zoo animals being kept in species-appropriate group sizes and having social needs met in mediating perceptions of welfare [19, 29, 32, 38, 54, 66, 72, 90, 93, 96, 107, 110, 111, 132], with social isolation being a primary factor of concern [32, 38, 66, 110, 111]. At the same time, one study identified that visitors also worry about overcrowding [29] and another showed that mixed species exhibits provoked concern if the mix included an animal perceived as dangerous [70]. Humans are generally considered a social species [140] and anthropomorphising the needs and feelings of animals is common [141]. Perceptions that “*no animal should be alone”* [66] are therefore unsurprising, but further research is required to identify what factors are influential to appropriate group size perceptions, particularly for species considered more solitary in nature.

##### Condition of the enclosure

The reviewed literature was consistent that zoo visitors expect enclosures to be clean and well maintained [16, 29, 32, 34, 40, 47, 64, 68, 80, 88, 98, 111, 125, 126, 129, 142], free from faeces [64, 70], safe [16, 70, 111, 126], and contain appropriate lighting [70, 94] and ventilation [70]. Additionally, newer facilities were perceived as more beneficial to welfare over older facilities [11, 66, 106].

##### Diet

Beliefs that zoo animals had access to adequate and appropriate diets and clean water were consistently associated with more positive welfare perceptions amongst visitors [11, 16, 19, 32, 39, 40, 43, 44, 54, 59, 68, 70, 90, 93, 96, 100, 101, 111], though visitors’ conception of what constitutes “appropriate” is unclear. Carcass feeding was perceived as beneficial to welfare [100, 101, 138], though reasoning behind such perceptions is unclear, as natural behaviour display was not always considered an indicator of welfare [100].

##### Sensory stressors

Exhibit locations exposing animals to noise from visitors and infrastructure were perceived to reduce welfare [32, 50, 68, 120], though noise in general or noise created by visitors was also sometimes perceived as enriching [50, 79, 99]. Proximity to other animals also provoked welfare concern [32, 47], though it is not clear if concerns focused exclusively on prey species and the potential stress from the threat of predation, or if concerns extended to the welfare of predatory species who may become frustrated by an inability to perform hunting behaviours.

##### Temperature

Perceived thermal comfort was influential to welfare perceptions in some cases [16, 33, 40, 47], although not others [19, 120]. This factor may therefore be more relevant for animals known to originate from more extreme climates that differ from the climate of the zoo location. For instance, polar bears were perceived as “*very pitiful, they are sitting in the heat all the time”* [40]. Nonetheless, provision of facilities to protect from weather exposure was expected and perceived as beneficial to welfare [33, 38, 47, 51, 120].

#### Welfare interpretation

Finally, several studies have suggested that interpretation regarding zoo animal welfare may be beneficial to improving visitor perceptions of welfare [2, 16, 19, 40, 75, 77, 95, 100, 104], but results are not consistent across studies. For example, perceptions of welfare seem to improve when interpretation includes information about the institution’s welfare accreditation, though improvement was lower in participants who initially perceived animals to be experiencing poor welfare [19]. In contrast, interpretation containing information regarding the choices provided to animals did not appear to impact visitor perceptions of welfare [75]. Similarly, several articles found that interpretation regarding care practices improved welfare perceptions [64, 69, 76, 77, 105], though only minor impacts were observed elsewhere [54], and some findings suggest that other variables such as animal activity may be more influential than interpretation [77]. Further exploration of the impacts of specific elements of interpretation may help clarify these inconsistencies.

### 3.6 Impact of welfare perceptions on visitor experience and behaviour

Negative animal welfare perceptions were commonly reported to have a negative impact on zoo visitor experience [10, 64, 98, 104, 107, 110, 131, 143] and satisfaction [16, 97, 125, 126, 131, 144, 145], though welfare perception was not always a contributing factor to visitor satisfaction [146]. Positive perceptions of welfare improved both visitor happiness (which in turn increased visitor enjoyment and improved experience) [103] and emotional experience/connection to the animals [16, 55, 75, 88, 109, 110]. In turn, positive emotional experiences and satisfaction were pivotal factors in the decision making of zoo visitors when considering revisiting [147], and emotional experience may also impact conservation learning [148]. In contrast, negative perceptions of welfare were reported to reduce the likelihood of visiting [2, 9, 56, 67, 106, 149–152], and donating to conservation efforts [2, 88, 103, 153].

Yet, the qualitative data presented in several of the reviewed studies suggest that negative perceptions of welfare may not always impact visit likelihood. For example, self-interest was a justification for captivity despite welfare concerns (see beliefs about captivity) and the desires of children to attend zoos was often given as an overriding factor in the decision to visit, despite conflicting negative welfare perceptions [9, 29, 67]. Cultural differences may also inform visit frequency despite welfare perceptions [149]. Negative perceptions of zoo animal welfare may therefore not actively prevent zoo visits, even by those who hold strongly negative beliefs about the ethics of zoos. The impacts of these conflicting feelings and actions, and how these in turn impact the behaviours of such ‘reluctant visitors’, should thus be more clearly addressed by future research.

In general, ‘non-visitors’ were considered to have lower perceptions of zoo animal welfare when compared to those defined as ‘zoo visitors,’ though welfare perceptions of both populations were influenced by similar factors like animal behaviour and enclosure size [18, 37, 106]. However, the term non-visitor seems to be used interchangeably to denote both people who choose not to visit zoos [37, 106], and those merely not on site at a zoo at the time of research [18, 37]. This may create ambiguous results, and therefore, having a definitive definition of non-visitors may be beneficial for meaningful comparisons. Godinez and Fernandez [3] encourage future inclusion of comparisons between visitors and non-visitors to provide a “*true control group*” to evaluate the efficacy of zoo-based education amongst these two populations. We propose defining ‘non-visitors’ as those who do not visit zoos. Sub-categories of non-visitors may consist of those who choose not to visit on animal welfare grounds, general lack of interest, cost, or other restricting factors. Each sub-category likely has differing opinions of zoos, which may impact perception research. ‘Zoo visitors’ may be defined as those who visit zoos and subcategorised as on or off site.

### 3.7 Summary

The objectives of this systematic review were two-fold: first, to identify the zoo attributes that influence visitor perceptions of zoo animal welfare, and second, to characterise how positive or negative perception of animal welfare impacts visitor attitudes, experiences, and behaviour. A total of 115 articles were reviewed and ultimately nine themes (3 human-level, 3 animal-level, and 3 environmental-level) described to capture attributes influencing zoo animal welfare perceptions (Obj. 1), with one theme (impact of welfare perceptions on visitor experience and behaviour) summarising how the valence of welfare perceptions consistently influenced visitor experiences and behaviour (e.g., negative perceptions contributing to negative experiences and vs versa, Obj. 2). Links between themes were evident, highlighting how and why each attribute was influential to welfare perceptions, which in turn have the potential to impact visitor attitudes, experiences, and behaviours.

Links between themes also provided insight into how zoos may potentially improve their communication with visitors regarding zoo animal welfare, but the theme *welfare interpretation* highlighted a need for further research to identify effective strategies to engage visitors with welfare related interpretation. As negative perceptions can result in visitor behaviours which may diminish the ability of zoos to achieve organisational goals such as conservation, education, and entertainment, establishing how zoos may effectively communicate with visitors regarding zoo animal welfare is important. However, of primary importance is the impact visitor perceptions may have on zoo animal welfare, as in the case of inappropriate visitor behaviours which may negatively impact animals in addition to other visitors’ experiences. Moreover, a better understanding of visitor perceptions may help highlight areas where zoos may benefit from improving. For example, the themes *condition of the enclosure* and *apparent health status* highlighted factors which (if perceived accurately by visitors) may detrimentally impact the welfare of captive zoo animals if not adequately addressed by zoos. More research is required to ascertain why visitors are reaching judgements of some themes, such as *enclosure size*, *group size,* and *animal behaviour,* to inform effective interpretation development.

### 3.8 Limitations

We acknowledge that studies meeting inclusion criteria may have been inadvertently missed despite our adherence to the PRISMA guidelines. Furthermore, the replicability of searches performed using Google Scholar can be impacted by factors such as location, and EBSCOhost results can vary dependant on institutional subscriptions [154]. Nonetheless, measures were taken to limit negative impacts to this review’s methodology (e.g., search strategy assessment by an academic librarian, independent searches by two researchers, and utilisation of four search systems). The exclusion of grey literature and studies not available in the English language may have introduced result bias, although inclusion can reduce search reproducibility [155] and introduce translation errors [156]. Nonetheless, potential publication bias is a limitation of this review.

### 3.9 Future research

While studies included in this review have provided valuable insight into how visitors perceive animal welfare in zoos, several gaps (Table 2), or inconsistencies were identified. We conclude by sharing seven recommendations to guide future work in this area (Fig 2). These recommendations focus on improving consistency in measures used to gauge visitor perceptions, as follows:

*1. Define key terms.* Ambiguity regarding how studies define key terms (e.g. animal welfare, non-visitors) can hinder direct comparisons. Additionally, as disagreement regarding definitions is evident between researchers [157], the potential for variations between visitors is probable. Establishment of clear definitions may therefore be beneficial to future research.
*2. Consider the dimensionality of welfare conceptions in measurement of perceptions.* Variations between how welfare perceptions are measured (e.g. welfare as happiness, health, ability to display natural behaviours, quality of care, overall welfare) can hinder direct comparisons between studies. Additionally, visitors consider some measurement categories to be more important to welfare than others, and category hierarchy varies between visitors [16, 19, 54]. Researchers risk missing valuable perspectives or even misrepresenting their research population’s views by imposing their own conceptions of welfare onto the measurement instruments. The inclusion of multiple welfare measurements, in addition to open-ended questioning, may therefore be beneficial to future research.
*3. Consider connecting themes for confounding impacts.* Connected themes (i.e. attributes highlighting how/why other attributes were influential to welfare perceptions) should be considered as potential confounding factors when designing perception research. Exclusion of connected themes may provide an incomplete or inaccurate overview of perceptions, and therefore inclusion may increase result validity.
*4. Include emotional impacts.* Emotional response to influential factors appears to be intertwined with experiences and perceptions [16, 55, 75, 88, 103, 109]: inclusion of emotional measurements is therefore also suggested to obtain a more in-depth understanding of how perceptions may impact visitor attitudes, experiences, and behaviours.
*5. Include a range of facility types.* How zoos choose to identify themselves (e.g. zoo, safari park, marine park) can impact welfare perceptions [39, 43, 44], yet most research has taken place within facilities identified as zoos. Expanding research to encompass a range of facilities may therefore be beneficial as results may not easily translate between facilities.
*6. Include a range of geographical locations.* Cultural differences can impact welfare perceptions [43, 149], yet research in this area is limited in several geographic regions, notably North America It may therefore be beneficial for research to expand and explore perceptions in a wider range of geographical locations.
*7. Include a range of animal classes.* Much of the work in this area has focused on perceptions toward the welfare of mammals; however, these perceptions do not necessarily translate to other animal classes, as human empathy and compassion towards animals can be dependent on how phylogenetically close the species is to humans [158]. Future research may therefore benefit from including an increased range of animal classes to understand welfare perceptions across a range of contexts.

**Table 2.**
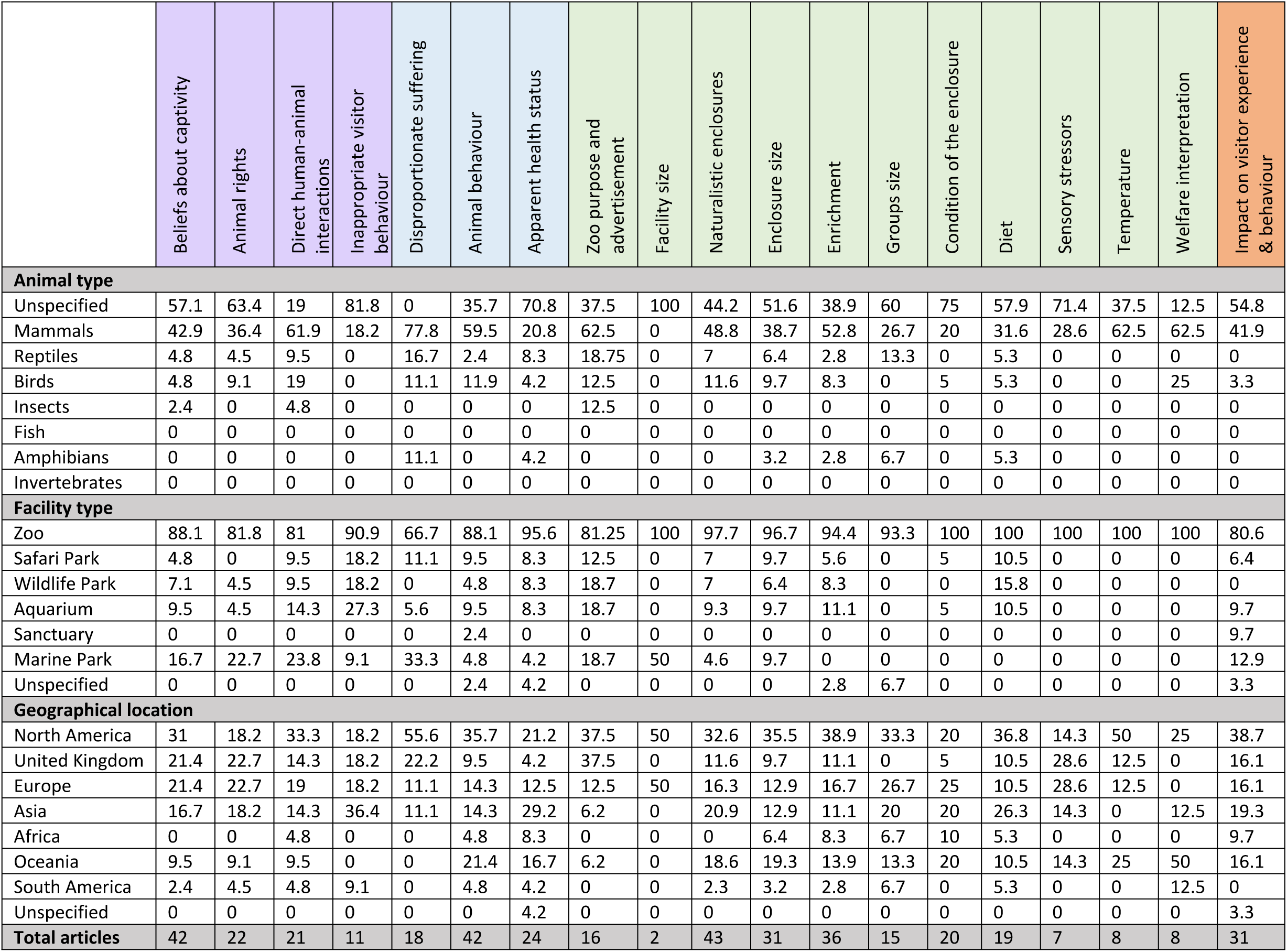
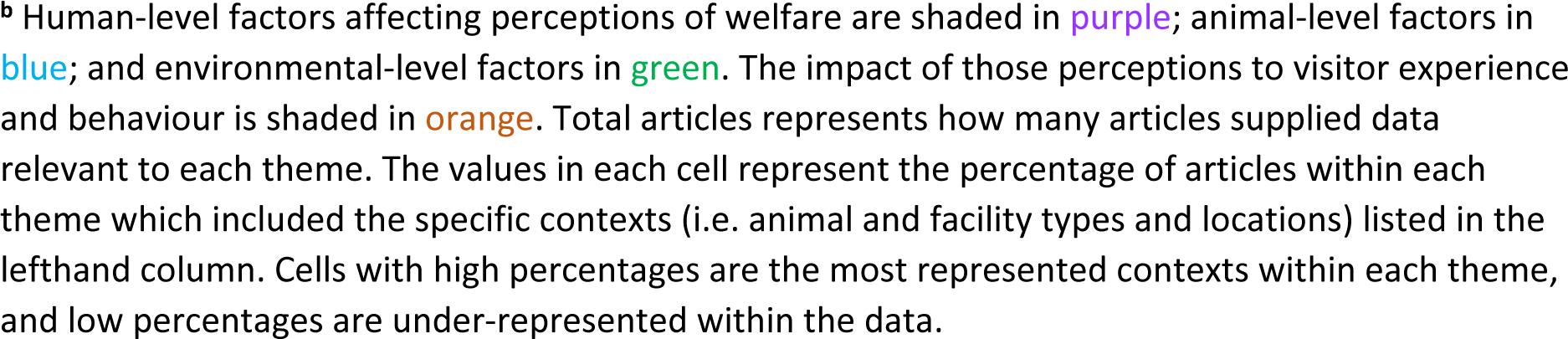
A summary of animal types, facility types, and geographical locations, which are currently represented or underrepresented in existing research for each theme. ^b^.

|  | Beliefs about captivity | Animal rights | Direct human-animal interactions | Inappropriate visitor behaviour | Disproportionate suffering | Animal behaviour | Apparent health status | Zoo purpose and advertisement | Facility size | Naturalistic enclosures | Enclosure size | Enrichment | Groups size | Condition of the enclosure | Diet | Sensory stressors | Temperature | Welfare interpretation | Impact on visitor experience & behaviour |
| --- | --- | --- | --- | --- | --- | --- | --- | --- | --- | --- | --- | --- | --- | --- | --- | --- | --- | --- | --- |
| Animal type |  |  |  |  |  |  |  |  |  |  |  |  |  |  |  |  |  |  |  |
| Unspecified | 57.1 | 63.4 | 19 | 81.8 | 0 | 35.7 | 70.8 | 37.5 | 100 | 44.2 | 51.6 | 38.9 | 60 | 75 | 57.9 | 71.4 | 37.5 | 12.5 | 54.8 |
| Mammals | 42.9 | 36.4 | 61.9 | 18.2 | 77.8 | 59.5 | 20.8 | 62.5 | 0 | 48.8 | 38.7 | 52.8 | 26.7 | 20 | 31.6 | 28.6 | 62.5 | 62.5 | 41.9 |
| Reptiles | 4.8 | 4.5 | 9.5 | 0 | 16.7 | 2.4 | 8.3 | 18.75 | 0 | 7 | 6.4 | 2.8 | 13.3 | 0 | 5.3 | 0 | 0 | 0 | 0 |
| Birds | 4.8 | 9.1 | 19 | 0 | 11.1 | 11.9 | 4.2 | 12.5 | 0 | 11.6 | 9.7 | 8.3 | 0 | 5 | 5.3 | 0 | 0 | 25 | 3.3 |
| Insects | 2.4 | 0 | 4.8 | 0 | 0 | 0 | 0 | 12.5 | 0 | 0 | 0 | 0 | 0 | 0 | 0 | 0 | 0 | 0 | 0 |
| Fish | 0 | 0 | 0 | 0 | 0 | 0 | 0 | 0 | 0 | 0 | 0 | 0 | 0 | 0 | 0 | 0 | 0 | 0 | 0 |
| Amphibians | 0 | 0 | 0 | 0 | 11.1 | 0 | 4.2 | 0 | 0 | 0 | 3.2 | 2.8 | 6.7 | 0 | 5.3 | 0 | 0 | 0 | 0 |
| Invertebrates | 0 | 0 | 0 | 0 | 0 | 0 | 0 | 0 | 0 | 0 | 0 | 0 | 0 | 0 | 0 | 0 | 0 | 0 | 0 |
| Facility type |  |  |  |  |  |  |  |  |  |  |  |  |  |  |  |  |  |  |  |
| Zoo | 88.1 | 81.8 | 81 | 90.9 | 66.7 | 88.1 | 95.6 | 81.25 | 100 | 97.7 | 96.7 | 94.4 | 93.3 | 100 | 100 | 100 | 100 | 100 | 80.6 |
| Safari Park | 4.8 | 0 | 9.5 | 18.2 | 11.1 | 9.5 | 8.3 | 12.5 | 0 | 7 | 9.7 | 5.6 | 0 | 5 | 10.5 | 0 | 0 | 0 | 6.4 |
| Wildlife Park | 7.1 | 4.5 | 9.5 | 18.2 | 0 | 4.8 | 8.3 | 18.7 | 0 | 7 | 6.4 | 8.3 | 0 | 0 | 15.8 | 0 | 0 | 0 | 0 |
| Aquarium | 9.5 | 4.5 | 14.3 | 27.3 | 5.6 | 9.5 | 8.3 | 18.7 | 0 | 9.3 | 9.7 | 11.1 | 0 | 5 | 10.5 | 0 | 0 | 0 | 9.7 |
| Sanctuary | 0 | 0 | 0 | 0 | 0 | 2.4 | 0 | 0 | 0 | 0 | 0 | 0 | 0 | 0 | 0 | 0 | 0 | 0 | 9.7 |
| Marine Park | 16.7 | 22.7 | 23.8 | 9.1 | 33.3 | 4.8 | 4.2 | 18.7 | 50 | 4.6 | 9.7 | 0 | 0 | 0 | 0 | 0 | 0 | 0 | 12.9 |
| Unspecified | 0 | 0 | 0 | 0 | 0 | 2.4 | 4.2 | 0 | 0 | 0 | 0 | 2.8 | 6.7 | 0 | 0 | 0 | 0 | 0 | 3.3 |
| Geographical location |  |  |  |  |  |  |  |  |  |  |  |  |  |  |  |  |  |  |  |
| North America | 31 | 18.2 | 33.3 | 18.2 | 55.6 | 35.7 | 21.2 | 37.5 | 50 | 32.6 | 35.5 | 38.9 | 33.3 | 20 | 36.8 | 14.3 | 50 | 25 | 38.7 |
| United Kingdom | 21.4 | 22.7 | 14.3 | 18.2 | 22.2 | 9.5 | 4.2 | 37.5 | 0 | 11.6 | 9.7 | 11.1 | 0 | 5 | 10.5 | 28.6 | 12.5 | 0 | 16.1 |
| Europe | 21.4 | 22.7 | 19 | 18.2 | 11.1 | 14.3 | 12.5 | 12.5 | 50 | 16.3 | 12.9 | 16.7 | 26.7 | 25 | 10.5 | 28.6 | 12.5 | 0 | 16.1 |
| Asia | 16.7 | 18.2 | 14.3 | 36.4 | 11.1 | 14.3 | 29.2 | 6.2 | 0 | 20.9 | 12.9 | 11.1 | 20 | 20 | 26.3 | 14.3 | 0 | 12.5 | 19.3 |
| Africa | 0 | 0 | 4.8 | 0 | 0 | 4.8 | 8.3 | 0 | 0 | 0 | 6.4 | 8.3 | 6.7 | 10 | 5.3 | 0 | 0 | 0 | 9.7 |
| Oceania | 9.5 | 9.1 | 9.5 | 0 | 0 | 21.4 | 16.7 | 6.2 | 0 | 18.6 | 19.3 | 13.9 | 13.3 | 20 | 10.5 | 14.3 | 25 | 50 | 16.1 |
| South America | 2.4 | 4.5 | 4.8 | 9.1 | 0 | 4.8 | 4.2 | 0 | 0 | 2.3 | 3.2 | 2.8 | 6.7 | 0 | 5.3 | 0 | 0 | 12.5 | 0 |
| Unspecified | 0 | 0 | 0 | 0 | 0 | 0 | 4.2 | 0 | 0 | 0 | 0 | 0 | 0 | 0 | 0 | 0 | 0 | 0 | 3.3 |
| Total articles | 42 | 22 | 21 | 11 | 18 | 42 | 24 | 16 | 2 | 43 | 31 | 36 | 15 | 20 | 19 | 7 | 8 | 8 | 31 |

<sup>b</sup> Human-level factors affecting perceptions of welfare are shaded in purple; animal-level factors in blue; and environmental-level factors in green. The impact of those perceptions to visitor experience and behaviour is shaded in orange. Total articles represents how many articles supplied data relevant to each theme. The values in each cell represent the percentage of articles within each theme which included the specific contexts (i.e. animal and facility types and locations) listed in the lefthand column. Cells with high percentages are the most represented contexts within each theme, and low percentages are under-represented within the data.

## 4.0 Conclusion

Key human, animal, and environmental factors were identified as influential to how visitors perceive zoo animal welfare, both positively and negatively, and these perceptions can subsequently impact visitor attitude, experience, and behaviour. Zoos must therefore consider how visitor concerns can be eased and positive perceptions enhanced, whilst primarily ensuring animals are indeed receiving high standards of care. Yet, several areas require further exploration to better understand what factors influence the perceptions of welfare in zoo animals, and research exploring how zoos may effectively communicate with visitors regarding animal welfare is currently limited. Further work addressing the gaps highlighted in existing research may benefit from increased consistency when gauging visitor perceptions of zoo animal welfare and may therefore benefit from the developed guidelines.

## Acknowledgements

With thanks to Andrew Cooke (Department of Life and Environmental Sciences, University of Lincoln) for his advice and guidance regarding table design and thanks to Carole Bee (University of Lincoln Library) for her advice and assessment of the search strategy.

## Supplementary materials

**S1. PRISMA checklist.**

**S2. Final search syntax.**

**S3. Pre-registered protocol.**

**S4. Articles excluded during screening.**

**S5. Mixed Methods Assessment Tool (MMAT) outcomes.**

**S6. Theme contribution (content/thematic) by each article included in the systematic review. S7. Theme overview (animal/facility/location/content/thematic).**

